# Intratelencephalic neurons in the medial prefrontal cortex mediate the acquisition of goal-directed actions

**DOI:** 10.64898/2026.07.02.736246

**Authors:** James N. Peak, Sophia Liang, Benjamin K. Lau, Karly M. Turner, Beatrice K. Leung, Bernard W. Balleine

## Abstract

The medial prefrontal cortex (mPFC) and its connections with the posterior dorsomedial striatum are implicated in goal-directed learning, but the specific mPFC cell types involved have not been clearly established. The current study investigated mPFC intratelencephalic (IT) and pyramidal tract (PT) neuron involvement in goal-directed learning. In Cre-driver mouse lines, we mapped bilaterally projecting IT neurons and unilaterally projecting PT neurons from mPFC to dorsal striatum and showed that chemogenetic inhibition of IT, but not PT neurons, attenuated goal-directed learning. We then demonstrated training induced elevations in pERK signaling in IT neurons, which were transient in superficial mPFC layers, and more persistent in deeper layers. This was associated with plasticity in deep layer IT neurons, reflected in a shift towards excitatory over inhibitory synaptic inputs. Together, these data suggest that goal-directed learning influences synaptic input and downstream plasticity markers in mPFC IT neurons, and this functionally contributes to goal-directed learning.

## Introduction

The acquisition of new goal-directed actions requires the encoding of action-outcome associations that, at a neural level, involves a prefronto-striatal circuit extending between the dorsomedial prefrontal cortex (mPFC) and the posterior dorsomedial striatum (pDMS) (Balleine, 2019; Balleine & O’Doherty, 2010). Within this corticostriatal circuit, the mPFC has been argued to play a specialized role as a short-term working memory, allowing the consolidation of specific action-outcome relationships for longer-term encoding in the pDMS (Balleine & Dickinson, 1998; Corbit & Balleine, 2003; Ostlund & Balleine, 2005; Stewart & Plenz, 2006; Choi et al., 2023). Thus, goal-directed learning has been found to increase the expression of cellular activity markers (e.g., phosphorylated extracellular signal regulated kinase (pERK)) in the mPFC, persisting for at least 1-hr after training has ceased (Hart & Balleine, 2016), activity that is required for the consolidation of action-outcome associations: blocking ERK phosphorylation immediately after training abolishes both plasticity in the pDMS and goal-directed learning (Hart & Balleine, 2016). Nevertheless, the subsequent retrieval of these associations appears not to depend on the mPFC (Ostlund & Balleine, 2005; Tran-Tu-Yen et al, 2009; Hart & Balleine, 2016), with considerable evidence instead implicating the pDMS in the longer-term maintenance and retrieval of action-outcome learning (Hart et al., 2014; Peak et al., 2020; Balleine et al., 2021; Choi et al., 2023).

While the above findings suggest that goal-directed learning is associated with learning-induced cellular plasticity in the mPFC, the specific cell types involved have yet to be confirmed. The rodent prefrontal cortex receives many long-range excitatory inputs (Hunnicutt et al., 2016, Hintiryan et al., 2016) and sends diverse outputs, primarily from deeper cortical layers (Tudi et al., 2024). In the context of goal-directed action, mPFC outputs to the pDMS are of particular interest and have been reported to be mediated by two distinct subclasses of pyramidal neurons - intratelencephalic (IT) and pyramidal tract (PT) neurons (Shepherd, 2013). IT neurons are present across cortical layers 2-6 and project bilaterally to other cortical or cortical-like regions, including the contralateral cortex, basolateral amygdala, claustrum, and additionally the striatum. By contrast, PT neurons are confined to layer 5b and project solely ipsilaterally to extratelencephalic structures, such as the thalamus, superior colliculus and the brainstem, whilst also sending collaterals to the striatum (Anastasiades & Carter, 2021). Therefore, while IT and PT neurons typically project to distinct regions throughout the brain, they have overlapping inputs into the striatum.

Whereas the anatomical distinctions between IT and PT cortical neurons have been well characterized, the functional contribution of these cell types to goal-directed action are relatively poorly understood. Recent evidence suggests a role of the IT pathway in goal-directed learning: after encoding new action-outcome associations, stimulating the cortico-striatal projections between the mPFC and pDMS in one hemisphere revealed changes in plasticity in pDMS principal neurons both ipsilaterally and contralaterally (Fisher et al., 2020). Furthermore, selectively silencing the contralateral projection from the mPFC to pDMS blocked the encoding of specific action-outcome associations during the acquisition of goal-directed actions (Hart et al., 2018a; 2018b). Nevertheless, to date the involvement of IT neurons in action-outcome learning has not been directly confirmed, nor has evidence of plasticity associated with such learning been reported in IT neurons.

The development of bacterial artificial chromosome (BAC)-Cre-Recombinase driver lines has enabled the investigation of cortical output neurons at a population level (Gerfen et al., 2013), providing the opportunity for a detailed assessment of IT neuron activity in goal-directed learning. Here, we used the KH288-Cre driver line, which selectively expresses Cre-Recombinase in IT neurons and the Sim1-KJ18-Cre driver line, which selectively expresses Cre-Recombinase in PT neurons, to address this question. Using pharmacogenetic manipulations, we found evidence that IT but not PT neurons are functionally required for the encoding of action-outcome associations during the acquisition of new goal-directed actions. Using fluorescence microscopy and electrophysiological recording experiments, we found evidence suggesting that the functional role of IT neurons is underpinned by elevated intracellular signaling during early learning, and a sustained longer lasting change in the synaptic input onto IT neurons favoring excitation.

## Results

### Tracing the output of medial prefrontal cortical IT and PT neurons to the striatum

We first sought to map the corticostriatal IT and PT neuron projections from the mPFC to the dorsomedial striatum, targeting, therefore, both the anterior region of the mPFC found previously to be important for the acquisition of goal-directed action and its striatal projection region generally regarded as the neural locus for goal-directed learning (Yin et al., 2005; Balleine, 2019). To achieve this, we took advantage of two transgenic BAC Cre-recombinase driver lines, Tg(Grp-cre)KH288GSat (“KH288-Cre”) and Tg(Sim1-Cre)KJ18GSat (“Sim1-Cre”), which respectively show specificity in corticostriatal IT and PT neurons (Gerfen et al., 2013). Prior work in other cortical regions confirms specificity of the targeted genes or cell types to excitatory neuron markers (Melzer et al., 2021, Marcassa et al., 2025) and projections specifically to intratelencephalic and pyramidal tract structures (Liu et al., 2024).

To label mPFC corticostriatal projections, we injected a Cre-dependent anterograde tracing virus, AAV2/5-hSyn-FLEX-mGFP-2A-Synaptophysin-mRuby (“Flex-mGFP-Synaptophysin-mRuby”) unilaterally into the mPFC of KH288-Cre and Sim1 -Cre mice (Figure 1A-D). This virus allows the visualization of axons through the expression of a green fluorophore (mGFP), and the visualization of downstream synaptic contacts through the expression of a red fluorophore (mRuby). We created density maps of synaptic connectivity (mRuby expression) in the dorsal striatum at different anteroposterior coordinates (Figure 1E). When ipsilateral and contralateral particle densities were analyzed, KH288-Cre mice (Figure 1F) showed no hemispheric difference (F(1,14)=0.90, p=0.36), consistent with bilateral striatal projections from mPFC IT neurons. In contrast, Sim1-Cre mice showed significant ipsilateral bias (F(1,14)=20.07, p=0.001), confirming predominantly ipsilateral striatal projections from mPFC PT neurons. These findings are consistent with the known bilateral and unilateral connectivity of IT and PT neurons, respectively. We found that in more anterior portions of the striatum, IT neuron connectivity was widespread, densely innervating large areas of both the ipsilateral and contralateral striatum. In more posterior portions of the striatum, the density of connections was isolated to the dorsal aspect of the striatum, which has previously been established as critical for instrumental conditioning (for review see Peak et al., 2019 and Balleine, 2019). By contrast, PT connectivity was exclusive to the ipsilateral striatum and, in both anterior and posterior regions, was sparsely expressed and localized to a more central area of the dorsal striatum.

**Figure 1.**
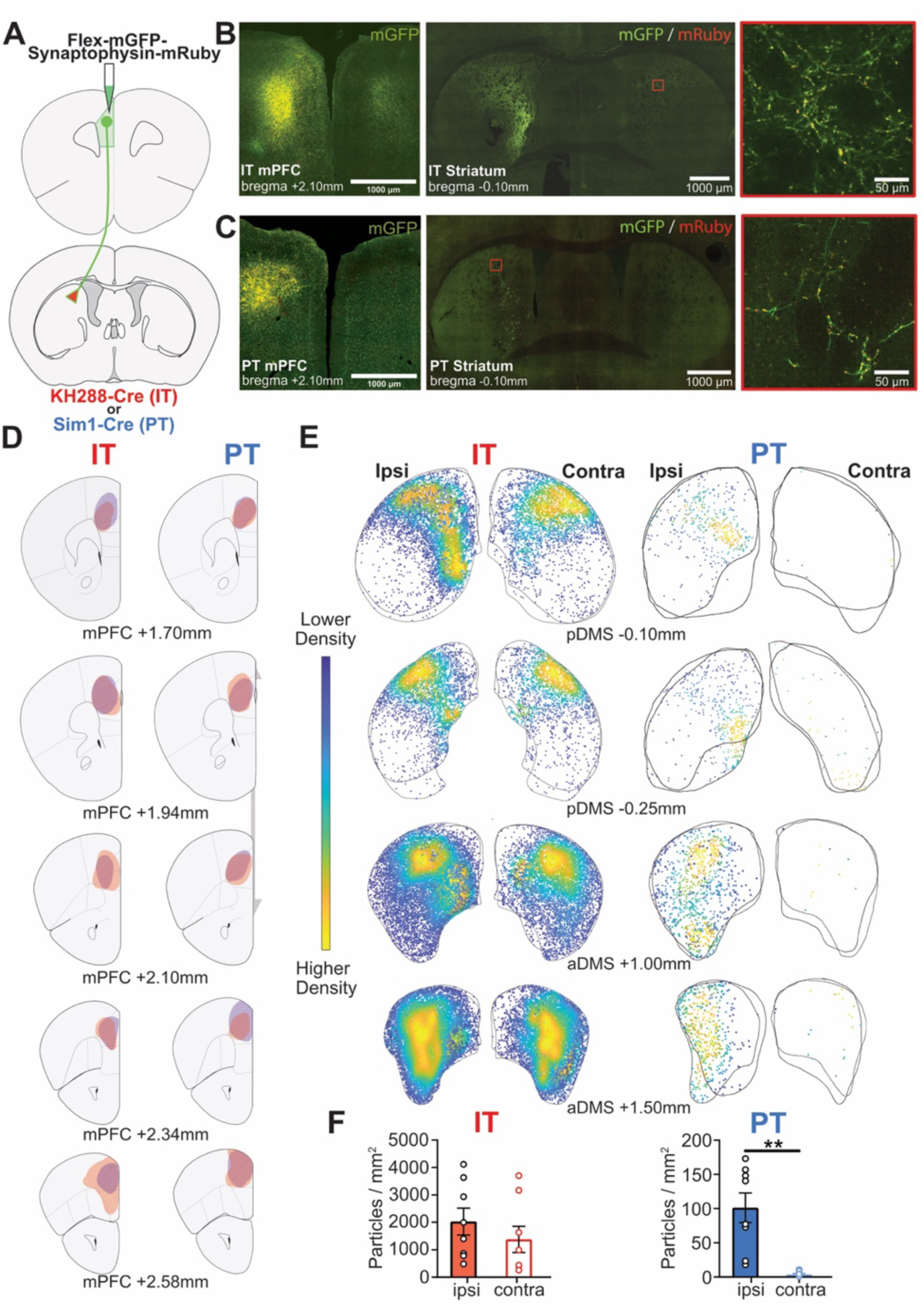
Projection mapping of IT neurons in KH288-Cre mice and PT neurons in Sim1-Cre mice. **(A)** Experimental schematic for circuit mapping of corticostriatal IT or PT neuron projections; KH288-Cre or Sim1-Cre mice received a unilateral injection of a Cre-dependent anterograde tracing virus, Flex-mGFP-Synaptophysin-mRuby targeted to the mPFC, which respectively labels cortical IT or PT neurons and their axons with mGFP (green), and their downstream synaptic contacts with mRuby (red). The localization and density of IT and PT neuron projections was mapped within the striatum. **(B)** Sample confocal image showing a unilateral injection site in the mPFC of a KH288-Cre mouse and labelling of IT neurons (left), mGFP and mRuby labelling in the ipsilateral and contralateral hemispheres of the striatum (middle), and a close up of the contralateral striatum from the location indicated by the red square in the middle image. **(C)** Sample confocal image showing a unilateral injection site in the mPFC of a Sim1-Cre mouse and labelling of PT neurons (left), mGFP and mRuby labelling in the ipsilateral striatum (middle), and a close up of the ipsilateral striatum from the location indicated by the red square in the middle image. **(D)** The extent of virus spread at the mPFC injection site for KH288-Cre (left) and Sim1-Cre (right) tracing animals mapped at five different anteroposterior coordinates. **(E)** Density mapping of mRuby puncta labelling at four different anteroposterior coordinates of the ipsilateral and contralateral striatum in KH288-Cre mice (N = 2) (left) and Sim1-Cre mice (right), overlayed on one another. **(F)** Bar graphs quantifying the density of mRuby puncta expression in the DMS of KH288-Cre (left) and Sim1-Cre mice (right). Density was calculated as the total number of particles per image divided by total area of each traced DMS to generate the mean number (±SEM) of particles per mm^2^ in the ipsilateral or contralateral striatum. All bars represent group means (±SEM). \*\**p*<0.01.

### Inhibition of mPFC IT neurons but not PT neurons impairs goal-directed learning

We next used a chemogenetic approach to assess the functional involvement of mPFC IT and PT neurons in goal-directed learning. KH288-Cre or Sim1-Cre mice received bilateral infusions of a Cre-dependent, inhibitory DREADD (Designer Receptor Exclusively Activated by Designer Drug) virus, AAV5-hSyn-DIO-hM4D(Gi)-mCherry (DIO-hM4D) virus or a control virus, AAV5-hSyn-DIO-mCherry (DIO-mCherry) into the mPFC (Figure 2A). For each experimental group (IT targeting or PT targeting), there were two control groups; one expressed DIO-mCherry in mPFC cortical neurons and received injections of Clozapine-N-Oxide (CNO) during training (“CNO control”), while the other expressed DIO-hM4D and received injections of vehicle (VEH) during training (“hM4D+VEH”). The third treatment group expressed DIO-hM4D and received injections of CNO (“hM4D+CNO”) during instrumental training to induce prolonged suppression of IT or PT neuronal activity. Schematics depicting viral spread at the injection site is presented in Figure 2B for KH288-Cre mice and confocal images showing DIO-hM4D expression in IT cells and counts of virus expression are presented in Figures 2C and 2D respectively. The same is shown for Sim1-Cre mice in Figures 2E-G. There were no differences in virus expression across groups in either experiment (IT or PT). We confirmed using *ex vivo* patch clamp electrophysiology that CNO produces functional inhibition of evoked firing in IT and PT neurons expressing DIO-hM4D, but not DIO-mcherry (Supplemental Figures S1A-F).

We then examined the effect of chemogenetic inhibition of IT and PT neurons on mice given relatively short instrumental training to learn two distinct action-outcome contingencies. Specifically, mice were first exposed to 4 days of drug-free magazine training for two distinct outcomes (O1 or O2), followed by 3 days of distinct A-O training where the same levers were paired with new, distinct outcomes (A1-O1 and A2-O2) (Figure 2A). Each distinct A-O training session occurred following an injection of CNO or VEH. It is worth noting that a small subset of animals in the IT neuron experiment received 5 days of distinct A-O training without pretraining (see methods for details). These animals were pooled with animals that received the protocol with pretraining, as there was no effect of protocol on press rates across training (highest F=3.01, Group hM4D+VEH) or test (highest F=1.63, Group hM4D+VEH) and there were no protocol x lever interactions or protocol x lever x time interactions in any group (highest F=1.91, Group CNO control) – see the Analyses section in the Methods. Following training, mice were tested for goal-directed action in a drug-free choice extinction test immediately after specific-satiety induced outcome devaluation. Goal-directed action is demonstrated by animals’ preferring the still valued lever, relative to the devalued lever, indicating an ability to incorporate current outcome value with knowledge of the specific A-O associations (Balleine & Dickinson, 1998).

**Figure 2.**
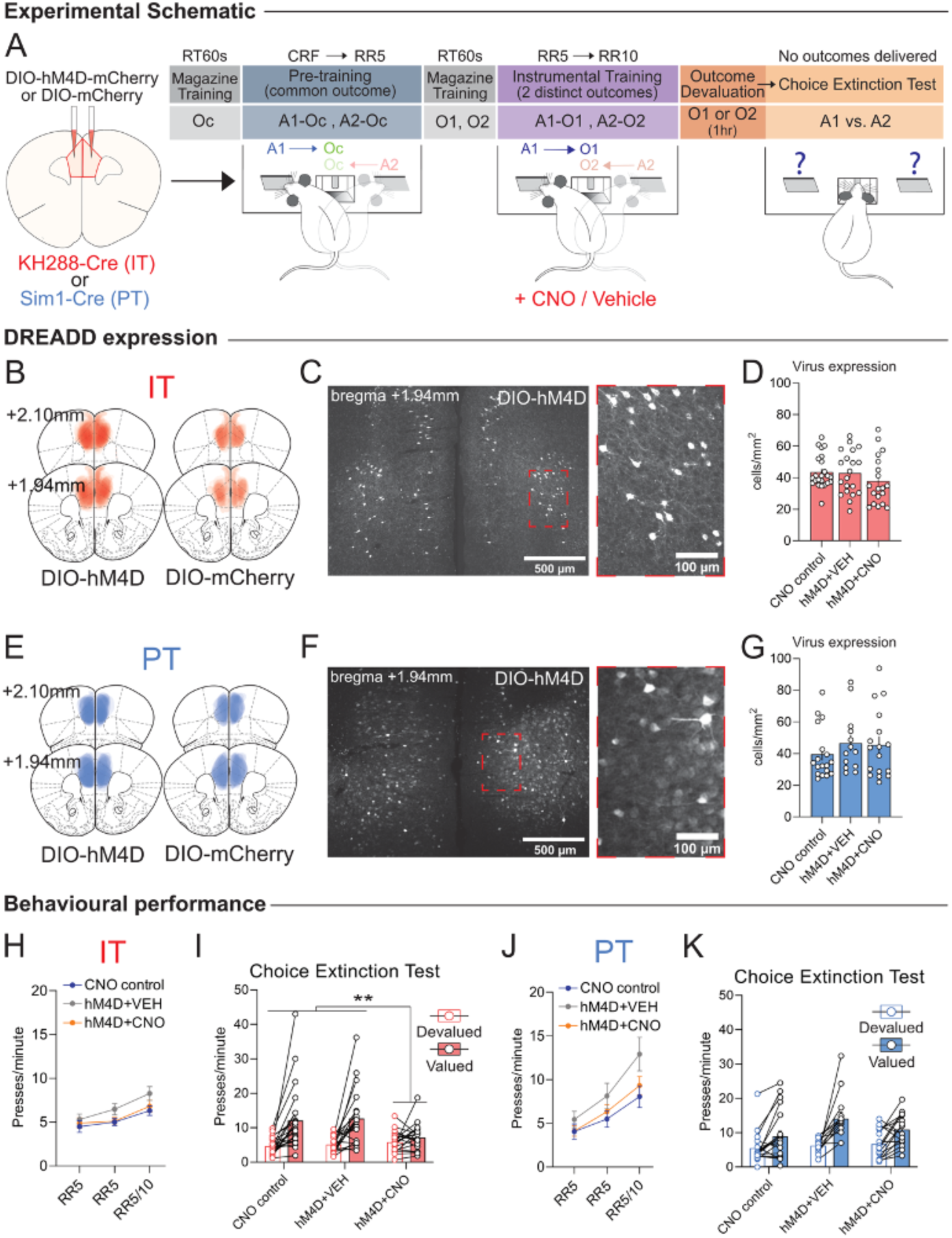
Inhibition of mPFC IT, but not PT neurons, impairs action-outcome learning. **(A)** Experimental schematic for chemogenetic inhibition experiment. Left, schematic depicting viral injections; KH288-Cre or Sim1-Cre mice received bilateral injections of a Cre-dependent inhibitory DREADDs virus, DIO-hM4D-mCherry or a Cre-dependent mCherry control virus (DIO-mCherry) targeted in the mPFC to infect IT or PT neurons. Right, schematic of the behavioural protocol; A1 and A2 indicate left or right lever press responses (counterbalanced across animals); Oc indicates a common outcome; O1 and O2 indicate two distinct food outcomes; CNO and Vehicle indicate days in which injections were administered; RT indicates random time; CRF indicates continuous reinforcement schedule; RR indicates random ratio schedule of reinforcement. **(B)** The extent of virus spread in the mPFC for all included KH288-Cre animals that received DIO-hM4D-mCherry (left) and DIO-mCherry (right), mapped at two different anteroposterior coordinates. **(C)** Confocal image showing DIO-hM4D expression in the mPFC of KH288-Cre mice. Inset: high power magnification of selected region highlighted in main figure. **(D)** Mean number (±SEM) of mCherry-expressing cells in the mPFC of KH288-Cre mice for each experimental group. **(E-G),** Same as B-D, but for PT targeting in Sim1-Cre mice. **(H)** Mean number (±SEM) of lever presses per min (averaged across both left and right levers) on each of the last three days of instrumental training for each group in KH288-Cre (IT) mice. **(I)** Mean press rate (±SEM) of the first half (2.5 mins) of the Choice Extinction Test (averaged across two days of testing) for each group in KH288-Cre (IT) mice. **(J-K)** Same as I-J but for PT targeting in Sim1-Cre mice. \**p*<0.05, \*\**p*<0.01, \*\*\**p*<0.001.

Training and test data for the IT inhibition experiment are presented in Figure 2H-I. Across the last 3 days of distinct A-O training, press rates increased (Figure 2H, linear trend, F(1,59)=63.30, *p*<0.001) and there were no significant differences between groups (highest F=3.42, hM4D+VEH vs CNO control), nor any group x day interactions (highest F=2.96, hM4D+VEH vs CNO control). However, it should be noted that there was a slight trend towards reduced responding in both the mCherry and hM4D groups receiving CNO. At test, the effects of a short A-O training period resulted in stronger extinction than expected. A comparison of changes in devalued and valued responding between the first and second half of the test showed a group (hM4D+CNO vs controls) x lever x time interaction (Bonferroni k=2, F(1,59)=12.18, *p*=0.001). This was driven by higher responding by control animals on the valued (vs. devalued) lever early in test, accompanied by rapid extinction on this lever, whereas animals with IT neuron inhibition during training maintained similar press rates on both levers at test. To confirm this, we conducted a further post-hoc contrast analysis on the first half of the test data (Figure 2I) and found no difference in press rates, averaged across levers (highest F=3.30, controls vs hM4D+CNO) but a main effect of devaluation (F(1,59)=32.39, *p*<0.001). Importantly, there was a group (control vs hM4D+CNO) x devaluation interaction (Bonferroni, k=2, F(1,59)=8.84, *p*=0.004) as control animals showed sensitivity to outcome devaluation but those which had IT neuron activity inhibited during test did not. Therefore, the inhibition of mPFC IT neurons during instrumental training attenuated the acquisition of action-outcome associations.

Behavioural data for the PT neuron inhibition experiment are presented in Figure 2J-K. Press rates increased across training day (Figure 2J, linear trend, F(1,45)=148.22, *p*<0.001) differently across groups; there was a group x day interaction (F(1,45)=9.38, *p*=0.004) and an overall group effect (highest F(1,45)=4.09, *p*=0.049, hM4D+VEH vs CNO control) as animals in the hM4D+VEH group had higher press rates. The two groups expressing the hM4D virus did not differ (F(1,45)=2.28, *p*=0.138). When examined for goal-directed action during the first half of the test (Figure 2K), there remained a main effect of devaluation (F(1,45)=40.57, *p*<0.001) and no group x lever interaction for the hM4D+CNO group when compared to the controls (Bonferroni, k=2, F(1,45)=1.34, *p*=0.253, controls vs hM4D+CNO). Further simple effect contrasts showed all groups exhibited sensitivity to outcome devaluation (lowest F(1,45)=8.21, *p=*0.006, hM4D+CNO). Therefore, whereas the inhibition of IT neurons in the mPFC during training attenuated goal-directed control, similar inhibition of PT neurons did not significantly affect the acquisition of action-outcome associations.

### Early goal-directed learning increases pERK expression in mPFC IT neurons

Given that the previous experiments revealed the functional necessity of mPFC IT neurons for the acquisition of action-outcome learning, we next investigated whether there was elevated activity and plasticity in these neurons by examining phosphorylated extracellular signal-regulated kinase (pERK) activation following goal-directed learning in the KH288-Cre mice. To examine co-localization with ERK, mPFC IT neurons were labelled with mCherry using the Cre-dependent fluorophore expressing virus, DIO-mCherry in KH288-Cre mice (Figure 3A). After 4 weeks recovery, all mice first underwent 4 days of magazine training. The following day, half of the mice received a single session of instrumental training in which pressing a lever (located to the left or right of the central food port) delivered a single outcome (grain pellets) on a continuous reinforcement (CRF) schedule, while the other half received yoked training in which outcomes were delivered at a rate determined by the master instrumentally trained mouse to which it was yoked (Figure 3B). This generated two groups: a master group for which lever pressing was associated with outcome delivery and a yoked control for which lever pressing and outcome delivery were unrelated.

**Figure 3.**
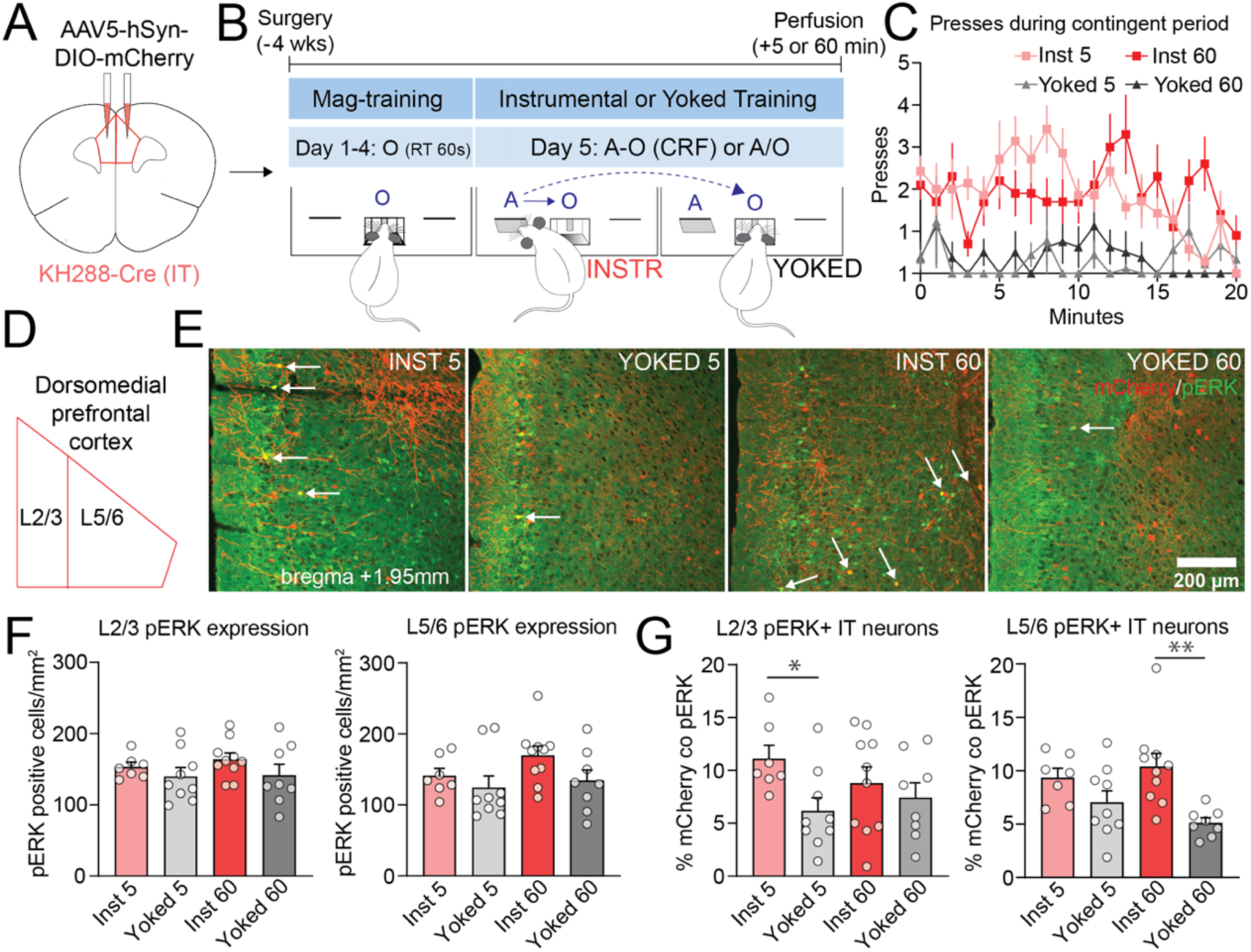
Early goal-directed learning increases pERK expression in mPFC IT neurons. **(A)** Schematic depicting viral injections for examining pERK expression in IT neurons; KH288-Cre mice received bilateral injections of a Cre-dependent fluorophore labelling virus, DIO-mCherry targeted in the mPFC to label IT neurons. **(B)** Schematic of the behavioural experimental protocol; instrumental animals were trained on a single lever (A) delivering grain pellets (O). Yoked animals had the lever present; however, outcome delivery was not contingent on a response and instead matched to a single instrumental animal. **(C)** Press rates (±SEM) for each group per minute during the contingent period of a single training session. **(D)** Region of Interests (ROIs) used for counting neurons in superficial (Layers 2/3) and deep (Layers 5/6) layers of the mPFC. **(E)** Example confocal images showing pERK expression (green), DIO-mCherry (red) and co-labelled cells (yellow - indicated by white arrows) at either 5 or 60 mins post-instrumental training in Yoked or Instrumental trained mice. **(F)** Mean number of pERK positive cells per mm^2^ in mPFC layers 2/3 (left) and layers 5/6 (right) per mouse in each group. **(G)** Percentage of labelled IT neurons that were co-labelled in layers 2/3 (left) and layers 5/6 (right) with pERK per mouse in each group. Data presented are means of five mPFC sections per mouse, averaged across hemispheres. All bars represent group means (±SEM). \**p*<0.05, \*\**p*<0.01.

To confine our analysis to early learning we used a criterion such that, when an instrumental animal pressed the lever twice within 60 seconds, they were deemed to have initiated learning. Meeting this criterion started a 20-min training period at the end of which half of each training group (Instrumental or Yoked) were perfused 5 mins post training (Inst 5, Yoked 5) to examine immediate pERK activation, whereas the other half were perfused 60 mins post-training (Inst 60, Yoked 60) to examine consolidation-relevant pERK activation. During the 20-min training period there was robust acquisition of the instrumental action in the master mice that was not apparent in yoked mice (Inst 5, Inst 60 vs. Yoked 5, Yoked 60, Figure 3C, F(1,30)=62.09, *p*<0.001).

Anatomically, the rodent mPFC is organized into 6 main layers (L), commonly subdivided into superficial L2/3 and deep L5/6. In general, L2/3 contains IT neurons that project to other cortical or cortical-like regions, whereas L5/6 contains a mixed group of IT neurons and PT neurons projecting to subcortical structures, including the ipsilateral and contralateral striatum (Anastasiades & Carter, 2021). When quantifying pERK expression alone in L2/3 and L5/6 IT neurons following instrumental or yoked training (Figure 3D), we found strong labelling of pERK in all groups (Figure 3E) with no main effect of instrumental training on overall pERK expression (Figure 3F) in mPFC L2/3 (F(1,30)=2.59, *p*=0.118) or L5/6 (F(1,30)=3.44, *p*=0.073). However, when analysing the percentage of the total labelled IT neurons that co-expressed pERK (Figure 3G), we found a significant effect of training type (Instrumental versus Yoked) in L2/3 (F(1,30)=5.07, *p*=0.032) and L5/6 (F(1,30)=13.54, *p*=0.001). Follow-up orthogonal contrasts compared each instrumental group with its matched yoked control (Inst 5 vs. Yoked 5 and Inst 60 vs. Yoked 60) and showed that instrumental training increased IT neuron pERK activity in L2/3 after 5 minutes (F(1,30)=5.80, *p*=0.022) and in L5/6 after 60 minutes (F(1,30)=13.94, *p*=0.001). Therefore, relative to yoked controls, instrumental training initially elevated pERK signaling in superficial mPFC IT neurons, followed by a delayed, or sustained elevation of pERK in deeper mPFC IT neurons.

### Goal-directed learning shifts the balance of IT neuron input to excitation over inhibition

Having established that early instrumental learning elevates pERK signaling, a marker of neuronal plasticity in IT neurons, we then assessed if this was associated with longer lasting changes in synaptic transmission. Similar to the previous experiment, mPFC IT neurons in KH288-Cre mice were fluorescently labelled with DIO-mCherry and mice were trained on the same instrumental training paradigm (Figure 4A). As expected, Instrumental but not Yoked control trained animals readily acquired the instrumental response (Figure 4B; F(1,18)=158.59, *p*<0.001).

Four hours following instrumental or yoked training or under behaviorally naive conditions, we used *ex vivo* patch-clamp electrophysiology to record spontaneous excitatory glutamatergic and inhibitory GABAergic synaptic transmission onto identified mPFC IT neurons (Figure 4C). In line with previously defined cortical layer boundaries (Anastasiades & Carter, 2021), recorded IT neurons were classified as L2/3 or L5/6 based on their measured distance from the hemispheric midline. The mean distance of recorded cells did not differ between groups (Figure 4D, Fs<2.6). We compared IT neuron recordings from Instrumentally trained animals against those from the Yoked + Naive control groups. In L2/3 mPFC IT neurons (Figure 4E), instrumental training produced no discernible change in either the rate or amplitude of miniature excitatory postsynaptic currents (mEPSCs) (Figure 4F; Fs<1.8) or inhibitory postsynaptic currents (mIPSCs) (Figure 4G, Fs<2.38) relative to control animals. To allow for a within-cell comparison of glutamatergic to GABAergic transmission, we further compared the rate of mEPSCs relative to mIPSCs within each individual neuron, calculating an excitation-inhibition (E-I) ratio. Overall, there was no difference in the E-I ratio between groups (Figure 4H, Fs<1).

**Figure 4.**
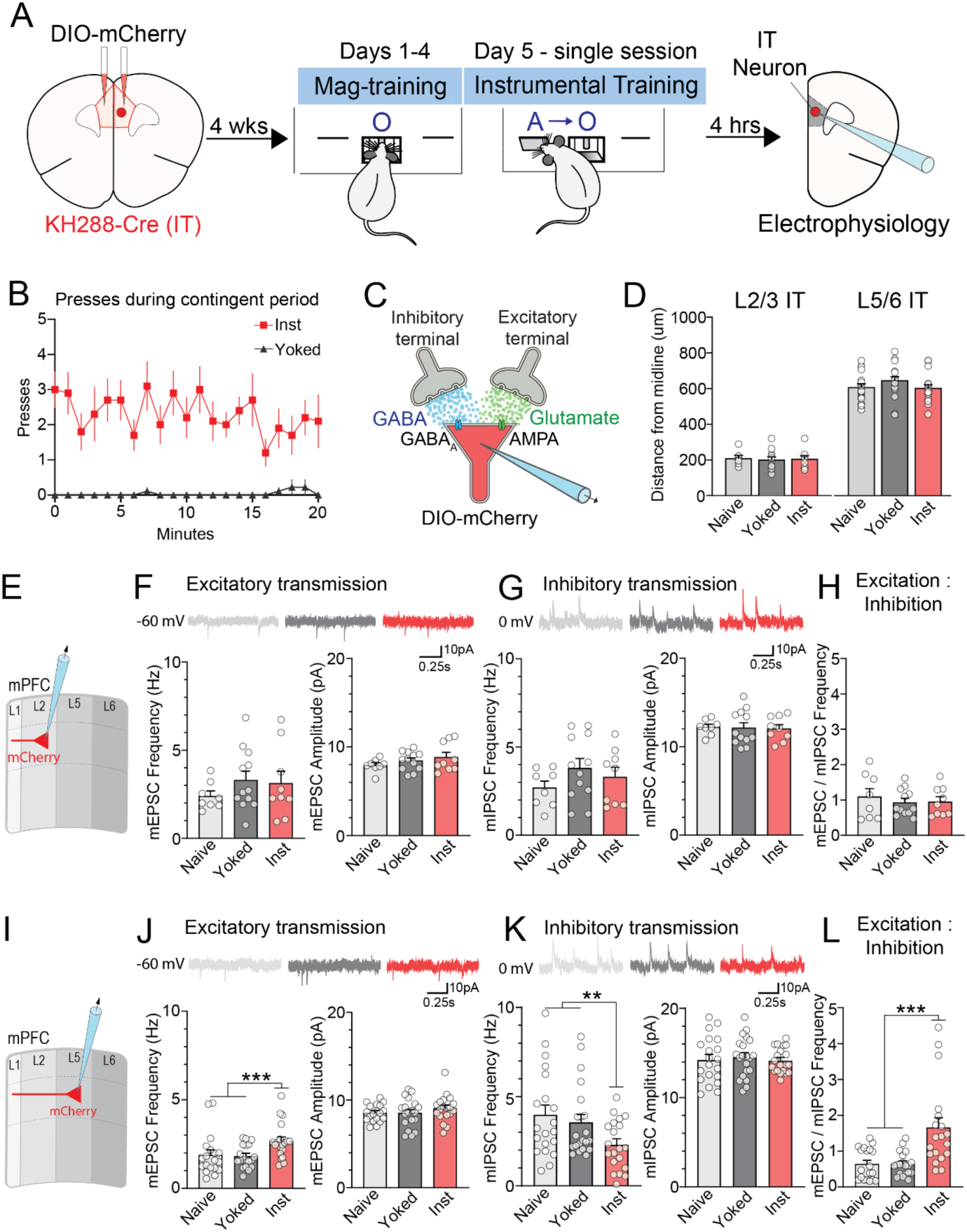
Early goal-directed learning induces a shift in the excitatory versus inhibitory input to IT neurons. **(A)** Schematic of the behavioural experimental protocol for examining learning-induced plasticity – KH288-Cre mice were bilaterally infused with DIO-mCherry virus in the mPFC. Four weeks post-surgery, mice underwent 4 days of magazine training, then received instrumental or yoked training over a single session on Day 5, followed 4 hrs later by ex-vivo electrophysiological recordings from mCherry+ IT neurons. **(B)** Press rates (±SEM) per minute during the contingent period of a single training session in the Yoked vs. Instrumental trained group. **(C)** Schematic of electrophysiological recording of synaptic transmission – ex-vivo patch-clamp recordings were made from mCherry+ IT neurons, specifically measuring spontaneous miniature excitatory or inhibitory postsynaptic currents (mEPSCs and mIPSCs), which reflect quantal glutamatergic and GABAergic synaptic transmission. **(D)** Distance from midline for each recorded IT neuron in Layers 2/3 (left) and Layers 5/6 (right) of the mPFC, in each group. **(E)** Schematic indicating the recording of Layer 2/3 mCherry+ IT neurons. **(F)** Upper traces show representative miniature EPSCs recorded in L2/3 IT neurons from a naive (grey), yoked (black) and instrumental (red) trained animal. Lower bar graphs quantify the frequency (left) and amplitude (right) of mEPSCs for each neuron per group. **(G)** Same as F, but for miniature IPSCs. **(H)** Bar graph quantifying the ratio of mEPSC to mIPSC rate in each L2/3 neuron per group. (**I–L)** Same as **E-H**, but for IT neurons in Layers 5/6. All bars represent group means (±SEM). \**p*<0.05, \*\**p*<0.01, *** *p*<0.001.

In L5/6 mPFC IT neurons (Figure 4I), instrumental training was associated with an increase in the rate (F(1,54)=9.03, *p*=0.004), but not amplitude (F<1.8) of mEPSCs (Figure 4J) relative to control animals, indicative of an enhancement in the presynaptic release of glutamate onto L5/6 IT neurons. In contrast, instrumental training was associated with a reduction in the rate (F(1,54)=6.77, *p*=0.012), but not amplitude (F<1) of mIPSCs (Figure 4K), suggesting a decrease in the presynaptic release of GABA onto L5/6 IT neurons. Importantly, the within-cell comparison of glutamatergic to GABAergic transmission revealed an overall enhancement of the E-I ratio compared to the control groups (Figure 4L; F(1,54)=23.87, *p*<0.001). Together, these results indicate that instrumental training is associated with a global, non-input specific shift in the balance of excitatory to inhibitory synaptic transmission in L5/6, but not L2/3 IT neurons.

## Discussion

The present series of experiments took advantage of transgenic Cre driver mouse lines to specifically target IT and PT pyramidal output neurons of the mPFC during the acquisition of goal-directed actions. We mapped the distribution of these distinct projection cell types and assessed their functional role in the acquisition of goal-directed actions, then investigated the intracellular signaling and plasticity mechanisms involved. We found that mPFC IT neurons send dense, bilateral projections to the dorsal striatum, including the pDMS, while PT neurons send a more sparse, ipsilateral striatal projection. Using chemogenetic inhibition of neuronal activity, we showed that IT but not PT neurons are functionally required for goal-directed learning. Associated with this involvement of IT neurons, we provided evidence that pERK expression is transiently elevated in L2/3 IT neurons immediately after early learning but is persistently elevated in L5/6 IT neurons for at least 1hr after training during a consolidation period. We then further demonstrated that L5/6 IT neurons exhibited lasting and divergent changes in spontaneous excitatory and inhibitory synaptic transmission, consistent with enhanced baseline excitability of this cell type following early instrumental learning.

### Projection-specific function: Goal-directed learning requires IT, not PT, neuronal activity

Goal-directed actions are the expression of learned associations between actions and their specific consequences or outcomes and are sensitive to changes in the value of those outcomes. Previous work from our lab has shown that pathway-specific inhibition of bilaterally projecting corticostriatal neurons in the mPFC during training abolishes goal-directed learning (Hart et al., 2018b). Here, we extended these findings to attribute a specific functional role of IT neurons, but not PT neurons, in this form of learning. Mice with chemogenetic inhibition of IT neurons during training were less sensitive to outcome devaluation in a choice extinction test; a task which requires knowledge of previously encoded action-outcome associations to guide choice.

Our results above are consistent with recent evidence demonstrating that prefrontal IT neurons are involved in encoding and maintaining information about outcomes during the delay period of a working memory task (Bae et al., 2021). Thus, IT neurons may serve as the substrate for working memory, required for maintaining new action-outcome contingencies during the early acquisition of goal-directed actions. However, it is clear that the bilateral corticostriatal pathway also conveys a learning signal to the pDMS during such learning, where it is additionally required for the expression or recall of action-outcome contingencies (Peak et al., 2019). In this regard, IT neurons innervate both populations of spiny projection neurons (SPNs) in the dorsal striatum - direct pathway SPNs (dSPNs) and indirect pathway SPNs (iSPNs) (Fisher et al., 2020). However, some recent evidence suggests functional activity is specific to dSPNs (Choi et al., 2023) and, indeed, plasticity at dSPNs but not iSPNs has been found to mediate goal-directed learning, with iSPNs involved in updating previously encoded associations when reward contingencies change (Peak et al., 2020; Fisher et al., 2020; Matamales et al., 2020).

The available evidence points, therefore, to a model of corticostriatal function in goal-directed learning wherein mPFC IT neurons broadcast a bilateral excitatory signal encompassing action-outcome information to the pDMS, where these signals converge with tightly timed dopamine signals from the substantia nigra pars compacta (Parker et al., 2016; Lee et al., 2019; Hart et al., 2024), promoting plasticity at dSPNs for the encoding of specific action-outcome associations. Consistent with this model, our current results show that a significant portion of mPFC IT neurons send a bilateral projection which targets the pDMS, and this is associated with a learning-induced elevation in the intracellular signal pERK, a key marker of synaptic plasticity. Furthermore, we showed that there is a learning-induced shift in the balance of synaptic inputs in favor of glutamatergic over GABAergic transmission in L5/6 IT neurons. Together, this change appears well placed to modulate the downstream excitatory plasticity of bilateral corticostriatal input previously reported in dSPNs, but not iSPNs in the pDMS following similar learning (Shan et al., 2014; Fisher et al., 2020). Indeed, there is evidence suggesting that IT neurons have a biased input onto dSPNS over iSPNs in other cortical regions, which if conserved in the mPFC-pDMS pathway, may potentially contribute to this cell-type specific plasticity (Reiner et al., 2010).

In contrast to IT neurons, we found that mPFC PT neurons did not contribute to goal-directed learning: although chemogenetic inhibition of PT neurons was effective at the cellular level (cf. Fig S1), their in vivo inhibition during training did not impact sensitivity to outcome devaluation at test. Although previous Cre-related mapping in IT and PT lines found expression was limited to dorsal mPFC (Gerfen et al, 2013), this lack of effect was not due to insufficient DREADD expression, as both hM4D-mCherry and mCherry fluorophore were expressed at similar levels as the earlier IT neuron experiments (see Fig. 2D vs. 2G). However, there is evidence that, at least in the motor cortex, PT neurons play a role in movement direction (Heindorf et al., 2018; Li et al., 2015) and in movement suppression in situations of conflicting actions (Reiner et al., 2010). Thus, PT neurons may contribute to elements of action initiation and execution during goal-directed behaviour (Moberg et al., 2022). While PT neurons may be important for other task-relevant functions, we found no significant changes to action performance when mPFC PT neurons were inhibited. Nevertheless, the neuronal inhibition typically observed with hM4D DREADDs is often incomplete and residual activity may have been sufficient to maintain performance rates. It should be recalled that PT projections to the dorsomedial striatum were found to be sparse, and, indeed, they have also been recently reported to only sparsely project onto striatal SPNs, however, they appear to send a more biased input to cholinergic interneurons relative to SPNs, an opposing circuit motif to that observed in IT neurons (Morgenstern et al., 2022). The behavioral relevance of this is unknown; however, given the prior established role of CINs in updating plasticity (in combination with iSPNs; e.g., Matamales et al., 2020; Peak et al., 2020), the contribution of the mPFC PT input to the striatum may be limited to the updating of instrumental contingencies if and when they change.

### IT neuron activity in goal-directed learning: Layer specific changes

Our circuit mapping experiment indicates that the mPFC sends a dense projection to the striatum, with IT neuron terminals concentrated in the dorsal aspect of the posterior region of striatum. This is consistent with the previously established role of a PL-pDMS pathway in instrumental learning (Hart et al., 2018b). These corticostriatal projection neurons are predominantly located in L5 of the PFC, which in turn are canonically modulated by superficial layers (Douglas & Martin, 2004; Anastasiades & Carter, 2021). Prior research has demonstrated that both L2/3 and L5/6 layers of the mPFC exhibit sustained activity, up to 1hr following a single session of instrumental training (Hart et al., 2016); however, the cellular identity of these activated neurons was unknown. Here, we specifically examined mPFC IT neurons following findings that they were functionally required for goal-directed learning. Here we found that early learning elevated pERK signaling in L2/3 and L5/6 IT neurons. Interestingly, layer specific activity appeared to be related to the interval after training: pERK elevation in L2/3 IT neurons was largely driven by instrumental trained animals perfused 5 minutes after training, whereas pERK elevation in L5/6 neurons was largely driven by instrumentally trained mice perfused 1-hr after training, although considerable activity was observed in both layers regardless of perfusion time. Relative to yoked controls, the pattern of IT neuron activation supports the view that action-outcome learning induces an immediate activation of IT neurons in the more input-heavy superficial cortical layers with a delayed activation of IT neurons emerging in more output-heavy cortical layers as learning is consolidated.

This sustained activity pattern in cortical neurons during action-outcome learning and consolidation is consistent with the involvement of a short-term, working memory process (Miller et al., 2018). Within the rodent cortex, connectivity between superficial (input) and deeper (output) layers is often emphasized and, indeed, intracortical excitatory connections between IT neurons, between IT and PT neurons and bidirectionally between superficial and deeper layers has been documented (Anastasiadis et al., 2021). Thus, the anatomical arrangement of this local circuit supports the amplification of recurrent activity in IT neurons. During goal-directed learning, excitatory inputs relevant to the sensory, affective, and motor properties of goal-directed actions innervate mPFC neurons (Balleine, 2019). These inputs converge with a dopaminergic error signal from the ventral tegmental area (Naneix et al., 2009) that enhances the excitability of IT neurons via signaling at the dominantly expressed dopamine type 1 receptors (Xing et al., 2021; Moburg et al., 2022). Additionally, inputs from the mediodorsal thalamus preferentially target L2/3 IT cells to provide a feedback signal as part of the broader cortico-striatal-thalamo-cortical circuit (Collins et al., 2018), reintroducing activation into the local mPFC circuit in a manner likely to support consolidation. As the mPFC appears not to be required beyond this consolidation phase (Ostlund & Balleine, 2005), we have previously argued that learning signals are transferred to the downstream dorsomedial striatum, providing the corticostriatal signals necessary to induce the plasticity at striatal principal neurons underlying the integration and storage of action-outcome associations (Peak et al., 2019); a hypothesis for which there is now considerable direct evidence (Fisher et al., 2020; Peak et al., 2020; Shan et al., 2014).

### IT neuron synaptic plasticity: Learning promotes a shift in the balance of excitatory to inhibitory synaptic input onto PFC IT neurons

It is well established that ERK activation is linked to the downstream induction of various immediate early genes, leading to the protein synthesis and synaptic plasticity processes that underlie learning and memory (Thomas & Huganir, 2004). Here, our observation of increased pERK following instrumental training suggested that long-term changes in synaptic plasticity may be present in mPFC IT projection neurons. Indeed, we found that most L5/6 IT neurons exhibited a dual alteration in synaptic transmission, with an increase in spontaneous glutamatergic input and decrease in GABAergic input, together resulting in a shift in the excitatory-inhibitory balance of synaptic inputs onto deep layer IT projection neurons under baseline conditions. Although the precise functional contribution of such synaptic alterations is unknown, they were observed only in the Instrumental group and not in the Yoked control group. This indicates plasticity specific to action-outcome learning, rather than from exposure to reward or context. Given that a significant proportion of L5/6 IT neurons in the mPFC send projections to the dorsal striatum (Anastasiades & Carter, 2021), also confirmed experimentally in our transgenic IT-neuron specific mouse line, it is likely that this plasticity occurs in a portion of striatal-projecting IT neurons; however, we cannot rule out changes in deep layer IT neurons projecting to other telencephalic regions. Using a similar behavioural paradigm, we have previously demonstrated learning-induced plasticity of downstream mPFC inputs in the pDMS (Fisher et al., 2020). Thus, our current observation of enhanced excitatory-inhibitory balance onto mPFC IT projection neurons may suggest a potential cellular mechanism underlying downstream adaptive alterations in the corticostriatal circuit. At present, we only report learning-induced changes in baseline synaptic transmission, and it is unclear whether similar synaptic changes occur in response to evoked action potential-dependent activity. Furthermore, the source of the excitatory and inhibitory input mediating these synaptic changes in L5 IT neurons is unknown. Therefore, future work will need to examine plasticity changes in an input specific manner evoked from either local intracortical neurons, or extrinsic regions previously implicated in action-outcome learning, including the basolateral amygdala (Fisher et al., 2020), mediodorsal thalamus (Corbit et al., 2003; Alcaraz et al., 2018) and hippocampal intermediary regions (Bradfield et al. 2020). This will aid in elucidating the upstream circuits driving the recruitment of the corticostriatal pathway during the acquisition of goal-directed actions.

## Conclusions

Taken together, our results demonstrate that goal-directed learning is critically dependent on the activity of prefrontal cortical IT neurons, which may be mediated by elevations in intracellular signaling and downstream alterations of their synaptic input. Our study utilizes modern Cre-transgenic mouse lines to undertake a detailed assessment of prefrontal IT and PT projecting neuron types in the acquisition of goal-directed action learning. Our findings are consistent with previous experiments conducted in rats that have implicated bilaterally projecting corticostriatal neurons in goal-directed learning (Hart et al., 2016; Hart et al., 2018b; Fisher et al., 2020) and highlight the absence of a functional role of PT neurons in such learning. Our findings also highlight a learning-induced shift in the balance of excitatory to inhibitory synaptic input onto mPFC IT neurons, which may be a potential cellular mechanism underlying their recruitment in goal-directed learning. This latter finding should be further investigated because, although the functional requirement of corticostriatal neurons in goal-directed learning is well established, the long-range and local intracortical inputs that are engaged during such learning are not yet clear. Just as IT-specific transgenic mice have allowed us to establish specificity of the effects reported here, there remain many open questions and, by combining these mice with in-vivo imaging tools, the subtle dynamics of IT neuron activity in the different cortical layers and various stages of action-outcome learning may be further revealed.

### Limitations of the Study

Given the current study utilised chemogenetics to target corticostriatal IT and PT neurons via a transgenic approach, this may only partially suppress a subset of the total IT/PT neuronal population. Specifically, our inhibitory DREADD verification indicates that CNO produced approximately 50% suppression of evoked action potential firing in both IT and PT neurons. Furthermore, our KH288-Cre and Sim1-Cre transgenic lines respectively target a molecular subtype of IT and PT neurons, which only partially overlap with the overall IT/PT neuronal populations targeted by other available BAC-Cre transgenic lines (Gerfen et al., 2013). Whilst chemogenetic inhibition was sufficient to impair goal-directed learning in mPFC IT neurons, it did not completely abolish sensitivity to outcome devaluation. Furthermore, the same manipulation in mPFC PT neurons had no effect on devaluation sensitivity. It is possible that the incomplete inhibition via chemogenetics may have contributed to the residual outcome devaluation sensitivity in this experiment. Likewise, we cannot rule out whether a more complete inhibition of mPFC PT neurons would have impaired goal-directed learning. Further work examining complete inactivation of both populations of mPFC projection neurons usubng, for example, cell-type specific tetanus toxins and cellular specific lesioning approaches would help to confirm the causal role of these cell types in goal-directed learning.

## RESOURCE AVAILABILITY

### Lead contact

Further information and materials requests should be directed to the lead contact, Bernard Balleine.

### Materials availability

This study did not generate any new materials.

### Data and code availability

- All animal data reported in this paper will be made available upon request to the lead author
- This study did not generate any original code
- Any additional information required to reanalyse the data reported in this study is available from the lead contact upon request.

## Acknowledgements

The research reported in this manuscript was supported by grants from the Australian Research Council, DP200103401, and the National Health and Medical Research Council of Australia, GNT1165346. The authors thank Charles Gerfen at NIH for his help getting access to the KH288-Cre and Sim1-KJ18-Cre driver lines.

## Correspondence

Bernard Balleine, Decision Neuroscience Laboratory, School of Psychology, UNSW Sydney, Randwick NSW 2052, AUSTRALIA.

## Methods

### Subjects

Adult (2-6 months age) male and female Grp-KH288-Cre mice (KH288-Cre) (Gerfen et al., 2013) were used for all IT experiments in this study. Adult (2-6 months age) male and female Sim1-KJ18-Cre mice (Sim1-Cre) (Gerfen et al., 2013) were used for all PT experiments. Mice were originally obtained from the Mutant Mouse Resource and Research Center and were then bred within the Decision Neuroscience Laboratory at The University of New South Wales. All mice were healthy and experimentally naïve at the beginning of each experiment and were at least 8 weeks old prior to surgery. Mice were housed in transparent, plastic boxes with two to four mice per box in a climate-controlled colony room and maintained on a 12 hr light/dark cycle (lights on at 0700). All experiments were conducted within the light phase. Water and standard lab chow were available ad libitum prior to the start of experiments. All experiments conformed to the guidelines on the ethical use of animals maintained by the *Australian code for the care and use of animals for scientific purposes*, and all procedures were approved by the Animal Care and Ethics Committee at The University of New South Wales.

For the IT and PT neuron tracing experiment, subjects were 2 female KH288-Cre mice and 2 female Sim1-KJ18-Cre mice, respectively. For the pERK experiment, subjects were 39 KH288-Cre mice (19 males and 20 females). For the electrophysiology experiment, subjects were 30 KH288-Cre mice (12 males and 18 females). For the IT and PT DREADDs inhibition experiment, subjects were 86 KH288-Cre mice (48 males and 38 females) and 94 Sim1-Cre mice (54 males and 40 females), respectively.

### Surgical procedures

Mice received stereotaxic surgery conducted under isoflurane gas anaesthesia (0.8L/min; isoflurane at 5% during induction and 2% during maintenance). Mice were placed in a stereotaxic frame (Kopf instruments) and a subcutaneous injection of bupivacaine at the incision site administered, as well as a subcutaneous injection of meloxicam in the lower flank. An incision was made to expose the scalp and the head was adjusted to align bregma and lambda on the same horizontal plane. 1-2 small holes were made above the PL cortex using a frame-mounted trephine and a capillary micropipette connected to a nanoject (Drummond Scientific) was lowered into the brain for infusions of viruses. Following each infusion, the needle was left for at least 4 minutes to allow virus diffusion. Following the final infusion, the incision site was closed and sutured. Mice received an antibiotic (Duplocillin; 0.15ml/kg s.c.) and saline (1ml i.p.) injection and were given a minimum of 4 weeks recovery to allow for viral expression.

Surgical coordinates were pre-determined from pilot studies. Viruses were infused into the dorsal portion of the mPFC at following coordinates (mm from bregma): AP +1.95mm; ML ±0.35 to 0.38mm; DV –2.1mm.

For tracing experiments: IT or PT neurons were unilaterally labelled with the anterograde tracing virus Flex-mGFP-mRuby (AAV2/5-hSyn-FLEX-mGFP-2A-Synaptophysin-mRuby, Addgene Viral Prep #71760-AAV1, Beier et al., 2015) at a total volume of 150nl and a rate of 120nl/min.

For pERK activity and electrophysiological recording experiments: IT neurons were bilaterally labelled with DIO-mCherry (AAV5-hSyn-DIO-mCherry, Addgene Viral Prep #50459-AAV) at a total volume of 500nl and a rate of 120nl/min.

For chemogenetic inhibition experiments, IT or PT neurons were bilaterally infected with either DIO-mCherry, or DIO-hM4D (AAV5-hSyn-DIO-hM4D(Gi)-mCherry, Addgene Viral Prep #44362-AAV) at a total volume of 450nl and a rate of 120nl/min.

### Behavioral procedures

For behavioral experiments, training and testing were conducted in 24 Med-Associates operant chambers, individually housed in light and sound attenuating cabinets. Each chamber was fitted with a food magazine that was connected to two 20mg pellet (Bioserve Biotechnologies) dispensers as well as two external pumps fitted with syringes that delivered either 0.075ml of 20% sucrose solution (white sugar, Coles, Australia) or 0.075ml of 20% maltodextrin solution (Poly-Joule, Nutrica, Australia). An infrared detector was situated horizontally across the inside of the magazine to detect head entries.

The chambers were also fitted with two retractable levers, located on either side of the magazine. A house light (3W, 24V) was in a central position at the top of the wall opposite the food magazine and illuminated during all experimental stages, except during inter-trial intervals (ITI). Training and test sessions were pre-programmed and controlled by computers external to testing rooms using Med Associates software (Med-PC V), which also recorded experimental data from each session.

For tests of goal-directed behavior via sensory-specific satiety, a separate devaluation room was used that contained 24 individual, open top plastic boxes with stainless steel wire mesh lids. During devaluation, room lights were kept off and each individual chamber was fitted with either a glass petri dish for pellet devaluation, or a plastic drink bottle with sipper attachment for sucrose devaluation.

#### Drugs

For DREADDs experiments, CNO (RTI International) was used to activate the hM4Di receptors and was made on the morning of each required day, dissolved in 0.1% 5M HCl, and diluted with distilled water (dH_2_O). Vehicle consisted of 0.1% 5M HCl diluted with dH_2_O and pH matched to the CNO solution. CNO was administered i.p. at a dose of 3mg/kg (Matamales et al., 2020).

#### Food restriction

For behavioral experiments, mice underwent 3 days of food restriction before the onset of magazine training. During this time, mice received 1.5g of standard lab chow for the first two days, and this increased to 2g for the third day and remainder of experiment, adjusted dependent on mice weights across days. Their weight was monitored closely to ensure that their food restricted body weight did not fall below 85% of their free-feeding body weight.

#### Magazine training

Mice were given an initial 3-4 days of magazine training during which the to-be-trained food rewards (either 20mg grain pellets or 0.075ml of 20% polycose solution (cued with clicker presentation)) were delivered at a random interval schedule of 60s. For the DREADDs inhibition experiment that included pre-training, an additional 2 days of magazine training for the two new outcomes (20mg grain pellets and 0.075ml of 20% sucrose solution (cued with second dispenser)) delivered at a random interval schedule of 60s were given in between pre-training and instrumental training.

#### Instrumental training

For *pERK activity* and e*lectrophysiological recording* experiments, mice received instrumental or yoked training sessions following 4 days of magazine training. Sessions began with the illumination of the house light and the presentation of either the left or right lever (lever position counterbalanced within groups). Lever press responses were rewarded with the delivery of 20mg grain pellets on a CRF schedule. The structure of the task was such that instrumental trained mice were given a 60-minute session to become contingent (two presses within the space of 60s, excluding the first press of the session). At the point of contingency, a 20-minute timer commenced and during this period, mice could earn as many pellets as their press rates allowed. Yoked animals received pellet delivery that was time-matched to an instrumental animal.

For *chemogenetic inhibition* experiments, a subset of mice underwent instrumental pre-training prior to instrumental training, and the other subset of mice underwent instrumental training without pre-training. For *pre-training,* both the left and right levers were rewarded with 0.075ml of 20% polycose solution. Sessions began with the illumination of the house light and the presentation of either the left or right lever (order counterbalanced within groups) for a quota of 5 outcomes or a period of 10 minutes, whichever came first.

Following this, the levers retracted, and the house light turned off for 2.5 mins before the alternate lever was presented. This sequence was repeated, such that all mice received two presentations of each lever. Days 1 and 2 were delivered on a continuous reinforcement schedule (CRF; every lever press earned one reward). On Day 3, animals that had fulfilled the criteria (successful completion (5 outcomes) on at least one block of each lever) on CRF were moved onto random ratio 5 (RR5; presses were rewarded on average every 5 presses). Animals that did not fulfil criteria on Day 2 remained on CRF for Day 3. On Day 4, all animals received training on an RR5 schedule. Following this, mice underwent 2 days of magazine training for grain pellets and 20% sucrose solution and then received 3 days of distinct action-outcome training, 30 mins after an injection of either CNO or VEH. During these training sessions, the same levers were paired with two new outcomes (20mg grain pellets and 20% sucrose solution, order counterbalanced within groups), on an alternating lever paradigm, whereby each lever was presented for 15 outcomes or 15 mins (whichever came first) and separated by a 2.5 min ITI, during which the house light turned off and the levers retracted. Day 1 and Day 2 of action-outcome training was on an RR5 schedule and Day 3 on an RR10 schedule.

For the subset of animals that received *instrumental training with no pre-training*, animals received distinct action-outcome training following magazine training (grain pellets and 20% sucrose solution) for 5 days, each day following an injection of either CNO or VEH. Mice were trained on a similar alternating lever paradigm as those who received pre-training, however for mice that did not receive pre-training, each block (left or right lever, counterbalanced within groups) a single lever was presented for 10 outcomes or 15 mins, whichever came first. Day 1-2 of training was on a CRF schedule of reinforcement, Day 3-4 was on an RR5 schedule and Day 5 of training was on an RR10 schedule.

#### Outcome devaluation

All mice were habituated to devaluation chambers with two 1 hr pre-exposure sessions during the training phase of the experiment. On test days, mice were placed in devaluation chambers for 1 hr where they had ad libitum access to one of the previously earned outcomes (grain pellets or 20% sucrose solution). Mice were then immediately returned to the operant chambers for a choice extinction test. Devaluation was conducted across two consecutive days of choice testing, with one outcome devalued each day, order counterbalanced within groups.

#### Choice extinction test

The choice-extinction test was conducted drug free. Sessions began with the illumination of the house light, and the simultaneous presentation of both left and right levers. No outcomes were delivered during this session, which ended after 5 minutes.

### Immunofluorescence procedures

#### Transcardial perfusion and tissue sectioning

For tissue analysis, mice received an intraperitoneal injection of a lethal dose of pentobarbital (0.6ml of a 1:10 solution diluted in saline) and were perfused transcardially with 75ml of cold 4% paraformaldehyde (PFA) in 0.1M phosphate buffer (PB). Fixed brains were immediately removed and stored in PFA for a further 12-36 hr. Brains were sliced into 30 µm coronal sections in 0.1M Tris Buffer Saline (TBS) solution on a vibratome (LeicaVT1000s) and stored at -30°C in a cryoprotective solution.

#### Immunofluorescence protocol

For *anterograde tracing experiments*, mGFP and mRuby were imaged without the addition of fluorescent antibodies in IT mice. Free-floating sections containing the striatum were washed twice for 10 min each in 0.1M TBS solution and twice for 10 min each in 0.1M Tris Buffer (TB) solution. For PT mice, free-floating sections containing the prefrontal cortex and dorsomedial striatum were washed three times for 10 min each in 0.1M TBS solution. After, sections were incubated in 0.5% Triton X-100, 10% normal horse serum (NHS) in 0.1M TBS for 2hr before being incubated in a solution containing a rabbit anti-DSRed antibody (1:1000), chicken anti-GFP (1:1000), 2% NHS, 0.2% TritonX-100 diluted in TBS at 4°C for 48 hr. Following this, sections were rinsed three times in 0.1M TBS for 10 min each before being incubated in solution containing a donkey anti-rabbit Alexa-594 (1:1000) and a donkey anti-chicken Alexa-488 antibody (1:1000), 2% NHS, 0.2% TritonX-100 diluted in 0.1M TBS for 2 hr. Finally, sections were rinsed three times for 10 min in TBS. All sections were mounted onto glass slides using Vectashield mounting medium. Images were taken using a Spinning Disk Microscope (Andor Diskovery).

For immunofluorescence in the *pERK activity experiment,* free-floating sections containing the prefrontal cortex were washed three times for 10 min each in 0.1M TBS solution containing 2% Sodium Fluoride (NaF). Sections then underwent peroxidase treatment in TBS (NaF) with 10% methanol and 3% hydrogen peroxide for 5 min. Sections were again rinsed 3 times in TBS (NaF) before undergoing a membrane permeabilization treatment for 15 minutes in TBS (NaF) with 2% Triton X-100 for 15 min. Sections were again rinsed 3 times in TBS (NaF) and then transferred into a solution containing the primary antibodies, rabbit anti-pMAPK (1:500) and chicken anti-mCherry (1:1000), diluted in TBS(NaF) for 36 hr at 4°C. Sections were washed 3 times in TBS (NaF) and then transferred into a solution containing the secondary antibodies, donkey anti-rabbit AlexaFluor 488 (1:1000) and donkey anti-chicken AlexaFluor 594 (1:1000), diluted in TBS (NaF) for 2 hr at room temperature. Sections were then rinsed twice in TBS and twice in (TB) for 10 min and mounted onto glass slides using Vectashield mounting medium. Images were taken using a Spinning Disk Microscope (Andor Diskovery).

For *chemogenetic inhibition experiments*, IT neurons expressing DIO-mCherry and DIO-hM4D were imaged without the addition of fluorescent antibodies. Free-floating sections containing the prefrontal cortex were washed two times for 10 min each in 0.1M TBS solution and two times for 10 min each in 0.1M TB solution. For PT neurons expressing DIO-mCherry and DIO-hM4D, free-floating sections containing the prefrontal cortex were washed three times for 10 min each in 0.1M TBS solution. After, sections were incubated in 0.5% Triton X-100, 10% NHS in 0.1M TBS for 2hr before being incubated in a solution containing a rabbit anti-DSRed antibody (1:1000), 2% NHS, and 0.2% TritonX-100 diluted in TBS at 4°C for 48 hr. Following this, sections were rinsed three times in 0.1M TBS for 10 min each before being incubated in solution containing a donkey anti-rabbit Alexa-594 antibody (1:1000), 2% NHS, 0.2% TritonX-100 diluted in 0.1M TBS for 2 hr. Finally, sections were rinsed three times for 10 min in TBS. All sections were mounted onto glass slides using Vectashield mounting medium. Images were taken using a confocal microscope (Olympus BX61WI) or a Spinning Disk Microscope (Andor Diskovery).

### Electrophysiology

#### In-Vitro Brain Slice Preparation

4 hrs following instrumental lever-press training, mice were deeply anaesthetized with isoflurane and intracardially perfused with cooled N-methyl D-glucamine (NMDG) solution of composition (in mM): 93 NMDG, 2.5 KCl, 1.2 NaH_2_PO_4_.H_2_O, 30 NaHCO_3_, 20 HEPES, 25 D-glucose, 5 sodium ascorbate, 3 sodium pyruvate, 2 thiourea, 10 MgSO_4_.7H_2_O, 0.5 CaCl_2_.2H_2_O (∼ 4°C, 300 mOsm). Mice were subsequently decapitated and coronal sections (250 µm) containing the mPFC were cut in NMDG solution using a vibratome (VT1200, Leica Microsystems). Slices were allowed to recover in warm NMDG solution (34 °C) for ∼ 10-12 min before being transferred to a holding chamber containing artificial cerebrospinal fluid (aCSF) of composition (in mM): 126 NaCl, 2.5 KCl, 1.2 NaH_2_PO_4_, 1.2 MgCl_2_, 2.4 CaCl_2_, 25 NaHCO_3_, 11 Glucose (34 °C, ∼ 300 mOsm); equilibrated with carbogen (95% O_2_ / 5% CO_2_).

#### Patch-Clamp Electrophysiology

Prior to recording, brain slices were individually transferred to a bath chamber on an upright microscope (Olympus BX51WI) and continuously superfused with aCSF (2 mL.min^-1^, 34 °C).

Neurons were visualized with a 40X water-immersion objective via Dodt gradient contrast optics. Whole-cell patch-clamp recordings of fluorescent mCherry-labelled IT neurons were conducted using a MultiClamp 700B amplifier (Axon Instruments, Molecular Devices). All recordings were filtered at 2 kHz, digitized at 10 kHz and collected using Axograph software. For measurement of synaptic transmission, voltage-clamp recordings were conducted using an internal solution of the following composition (in mM): 137.5 CsCH_3_SO_3_, 8 CsCl, 10 HEPES, 0.25 EGTA, 7 Na_2_-phosphocreatine, 4 MgATP, 0.3 NaGTP (pH 7.2-7.3, ∼ 305 mOsm). To specifically compare excitatory versus inhibitory transmission, spontaneous miniature excitatory postsynaptic currents (mEPSCs) or inhibitory postsynaptic currents (mIPSCs) were electrically isolated within the same neuron by respectively recording at a holding potential of -60 mV or 0 mV, in the presence of the NMDA receptor antagonist, DL-AP5 (50 µM, Tocris) and sodium channel blocker, tetrodotoxin (0.3 µM). Each mEPSC and mIPSC recording was run for a duration of 300 s with input and series resistance (< 20 MΩ) routinely monitored before and after (excluded if > 20% change). Spontaneous synaptic events were detected using a sliding template algorithm in the Axograph Software. For measurement of neuronal firing, current-clamp recordings were conducted using an internal solution of the following composition (in mM): 130 K-Gluconate, 10 KCl, 10 HEPES, 4 MgATP, 0.3 NaGTP, 10 Na_2_-phosphocreatine (pH 7.2-7.3, ∼ 295 mOsm). Neurons were maintained at resting membrane potential and action potential firing was evoked every 20 secs via current injection (1 sec) through the patch electrode. In each neuron, the magnitude of current injection was adjusted to elicit between 5-25 action potentials over time.

### Analyses

All experiments and analyses which follow were conducted in a non-blinded fashion by the experimenter. All statistical tests used an alpha level of 0.05. For post-hoc analyses, a Bonferroni correction was applied throughout to adjust for multiple comparisons.

### Exclusions and group allocation

pERK activity experiment – 5 mice were excluded; 3 for low virus expression (<4 infected cells) and 2 animals (one pair, master and yoked) that did not acquire the instrumental response. This left 34 mice, of which 19 were males and 15 were females (Group Inst 5, n=7; Group Yoked 5, n=9; Group Inst 60, n=10; Group Yoked 60, n=8)

Electrophysiology recording experiment: 3 mice were excluded due to technical issues on the day of recording, leaving 27 animals, of which 11 were males and 16 were females (Group Instrumental, n=10; Group Yoked, n=9; Group Naïve, n=8).

IT DREADDS inhibition experiment: 24 mice were excluded; 19 for low virus spread (<20 infected cells), 1 for lateral virus spread and 3 that did not acquire the instrumental response. This left 62 mice for analysis, of which 30 were males and 32 were females (Group hM4D+CNO, n=19; Group hM4D+VEH, n=20; Group CNO control, n=23).

PT DREADDs inhibition experiment: 46 mice were excluded; 41 for low virus spread (<20 infected cells) and 5 that did not acquire the instrumental response. This left 48 mice for analysis, of which 33 were males and 15 were females (Group hM4D+CNO, n=17; Group hM4D+VEH, n=13; Group CNO control, n=18).

### Immunofluorescence analysis

All immunofluorescence analysis was conducted using image J software. Animals with misplaced injection sites were excluded. Density mapping of anterograde tracing was performed using ImageJ and Matlab. Statistical analyses were performed using PSY software. In all analyses, the per-comparison error rate was controlled at α=0.05.

#### Tracing analysis

Image acquisition and particle analysis was performed in line with methodology published previously (Ferguson et al., 2024). Single 20x magnification images of brain slices containing the striatum at different anteroposterior coordinates were obtained using a spinning disk confocal microscope (Andor Technology, Nikon) and analysed using ImageJ/Fiji software).

The freehand tool was used to trace the perimeter of the striatum in each image, and the coordinates (x, y) of the points defining the line obtained as well as the total area of each traced striatum (mm*2*). For a set of images, a threshold was set based on the pixel intensity of the synaptophysin (mRuby) fluorescence and a binary image generated. The “Analyze Particle” function in FIJI/Image J was used to detect fluorescent particles above this threshold in each binary image. This function locates the edge of an object within the image based on size and roundness, and determines the coordinates (x, y) of its centroid position. For each image, coordinates of the striatum perimeter and coordinates of all detected particles were exported in a tab-delimited spreadsheet format and imported into MATLAB. For each image, the “inpolygon” function was used to generate a line plot defined by the coordinates of the striatum and a scatter plot defined by synaptophysin particle centroid coordinates. The “densityplot” function was used to colour code each plotted particle relative to the spatial density of nearby plotted particles, in each individual image. Using the number of detected particles for each image and the area of each traced striatum, a particle density value was calculated (particles/mm*2*).

#### pERK activity experiment

To maintain consistency across slices, sections within a series were imaged the day following immunofluorescence staining and all images taken within an 8 hr period. Quantifications were performed manually for mPFC L2/3 and L5/6 by defining separate region of interests (ROIs) that encompassed each layer class. All cells that demonstrated pERK expression, mCherry expression, and those co-labelled were counted. Total cell counts were analyzed as a total number of cells, calculated as the mean number of cells per mm^2^, per hemisphere, averaged across five mPFC slices per mouse. IT neuron activity was calculated as a percentage of the total labelled pathway; cells that were positive for pERK and mCherry were expressed as a percentage of the total number of mCherry positive cells in that slice.

The two different layer classifications were analyzed separately. Orthogonal contrasts were used to test for a main effect of training (Inst versus Yoked) and time (5 mins versus 60 mins) and the interaction between the two. A separate set of orthogonal contrasts was performed to compare each instrumental training group (Inst 5 or Inst 60) with the paired yoked control group (Yoked 5 or Yoked 60).

#### Chemogenetic inhibition experiment

Counts of mCherry positive cells in the mPFC were conducted manually and total number of cells expressed as cells/mm^2^. Orthogonal contrasts were used to compare the number of cells expressing either DIO-hM4D or DIO-mCherry and the number of cells expressed DIO-hM4D between the two groups that received this virus.

#### Electrophysiological analysis

Data from electrophysiological recordings were analyzed using orthogonal contrasts, comparing recordings from IT cells in Group Instrumental against the two control groups; Group Yoked and Group Naive) and directly comparing between the two control groups. This analysis was performed separately for cell recordings obtained in L2/3 and L5/6 and included measures of distance from midline, mEPSC rate, mEPSC amplitude, mIPSC rate, mIPSC amplitude, and Excitation:Inhibition (E-I) ratio (mEPSC rate/mIPSC rate). Statistical analyses and graphs were generated using Graphpad Prism software. The per-comparison error rate was controlled at α=0.05.

#### Behavioral analysis

For behavioral experiments, data collection was performed using Med-PC software (Version V), extracted using MPC2XL (Version 1.4, Med Associates) and tabulated using Microsoft Excel. Statistical analyses were performed using PSY software and graphs were generated with GraphPad Prism 9. The per-comparison error rate was controlled at α=0.05. For post-hoc analyses, a Bonferroni correction was used to control the family-wise error rate at α=0.05.

#### pERK activity experiment and electrophysiological recording experiment

Data from the 20-minute contingent period was analyzed using orthogonal contrasts, testing for a main effect of group (instrumental versus yoked) on press rates during this period.

#### Chemogenetic inhibition experiment

Planned orthogonal contrasts were used for training data analyses. Training data was analyzed testing for a main effect of group (hM4D+CNO versus controls, hM4D+VEH versus CNO control) and a main effect of training day and any interactions. Due to the rapid extinction in responding, post hoc orthogonal contrasts were used to analyze choice extinction test data. Within-group interactions for lever (devalued versus valued) x time (first versus second half of test) were conducted for all groups. Between-group analyzes of the first half of test were conducted for a main effect of group, a main effect of lever, and any interactions (group x lever). The FWER was controlled at α=0.05 using Bonferroni correction.

## Supplemental Figures S1A-F

**Figure S1:**
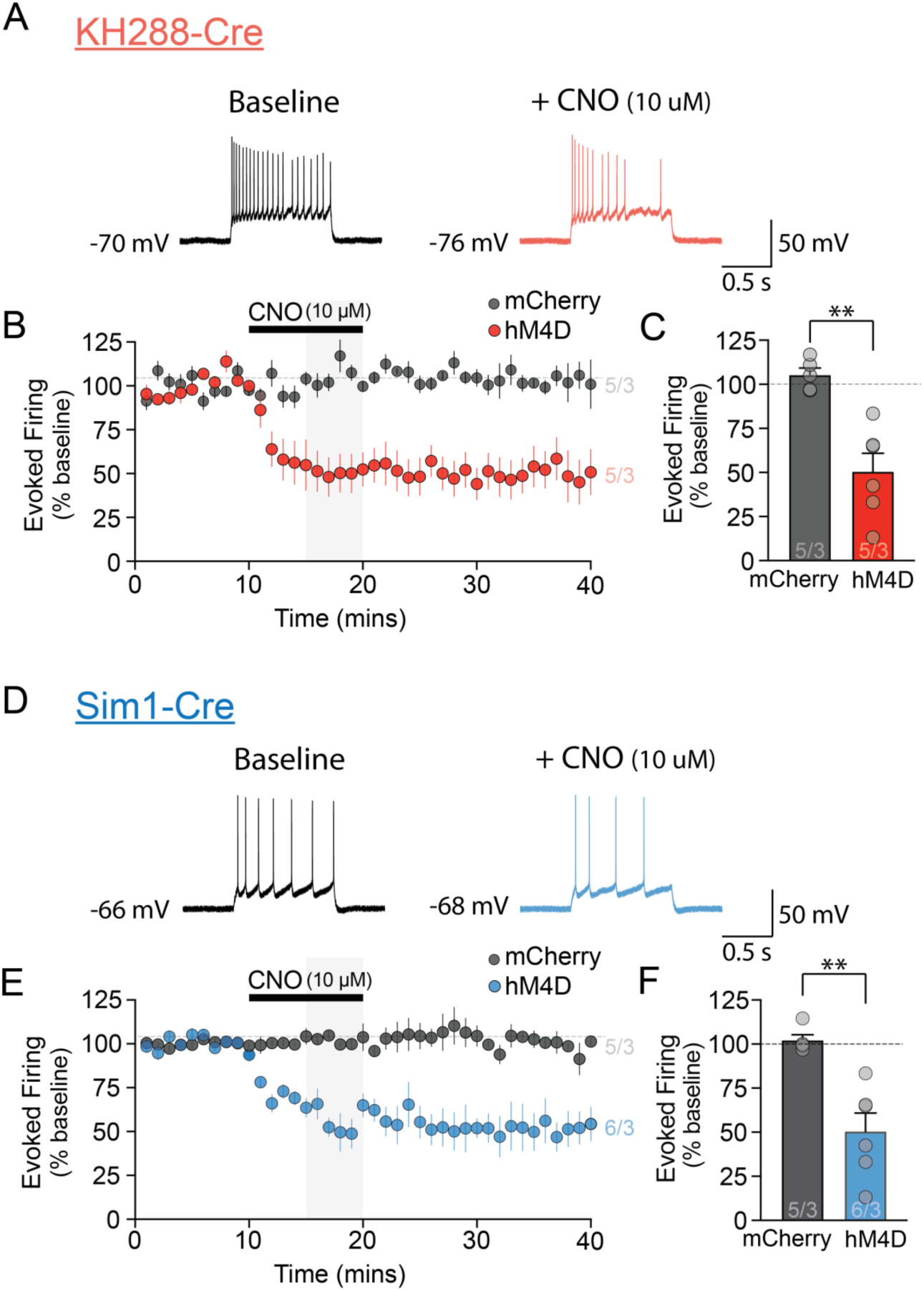
*ex vivo* validation of chemogenetic inhibition in IT and PT mPFC neurons. **(A)** Sample traces of evoked action potential firing before and during application of CNO (10 μM) in hM4D-expressing neurons of KH288-Cre mice. **(B)** Time plot of evoked firing before, during and after 10 min application of CNO in hM4D vs. mCherry expressing neurons in KH288-Cre mice. Firing is expressed as a % of baseline. Shaded area indicates the time period analysed in (C). **(C)** Bar graph quantifying the percentage inhibition by CNO (during 5-10 mins application) in KH288-Cre mice. **(D)** Sample traces of evoked action potential firing before and during application of CNO (10 μM) in hM4D-expressing neurons of Sim1-Cre mice. **(E)** Time plot of evoked firing before, during and after 10 min application of CNO in hM4D vs. mCherry expressing neurons in Sim1-Cre mice. Firing is expressed as a % of baseline. Shaded area indicates the time period analysed in (F). **(F)** Bar graph quantifying the percentage inhibition by CNO in Sim1-Cre mice. All bars represent group means (±SEM). \*\**p*<0.01.

